## Supplemental Information for "TGF-β1 activates neutrophil signaling and gene expression but not migration"

**Legends to Supplementary Figures**

**Figure S1: fMLF and CXCL1 stimulate pERK1/2.**

**(A)** Immunoblots of pERK1/2 and GAPDH in dHL-60 cells in response to fMLF or CXCL1 from 0 to 20 min. **(B)** Quantification of (A). ***P ≤ 0.0001, ****P ≤ 0.0001 when compared with time 0 (untreated (UT)) (one-way ANOVA with Dunnett's multiple comparisons test).

**Figure S2: Changes in the environment of dHL-60 cells affect gene expression.**

**(A)** Venn diagram depicting the number of changed genes in the IMDM and DMEM/F12 media conditions when each were compared to untreated control (t0). **(B, C)** Heat maps of Log2 Fold Change to mean time 0 expression in the (B) 'chemokine receptors bind chemokines (R-HSA-380108)' reactome pathway and the (C) 'neutrophil degranulation (R-HSA-6798695)' reactome pathway in dHL-60 cells either untreated (time 0) or treated with media controls (IMDM or DMEM/F12), TGF-β1, or M4 TCM for 30 min. Heat maps in (C) depict the 30 pathway genes with the largest average upregulation or the 30 genes with the largest downregulation relative to untreated.

**Supplementary Table 1** Significantly changed genes within specific reactome pathways.

| Reactome Pathway | Gene | Log2FC |
|---|---|---|
| Signaling by TGF-β family members | JUNB | 1.095 |
| | TGIF1 | 0.882 |
| | SMAD7 | 3.375 |
| | PAI1 | 1.724 |
| Class A1 (rhodopsin-like receptors) | ADRB1 | -1.645 |
| | CXCL3 | -1.705 |
| | EDN1 | -2.191 |
| | GPR35 | 0.989 |
| | S1PR2 | 3.137 |
| | GPR65 | -0.619 |
| | PTGER4 | -0.728 |
| | S1PR1 | -2.048 |
| Signaling by interleukins | VEGFA | 1.185 |
| | OSM | 4.279 |
| | JUNB | 1.095 |
| | CEBPD | -0.617 |
| | CISH | 1.036 |
| | SOCS1 | 1.464 |
| | NFKBIA | 0.822 |
| | CDKN1A | 1.592 |
| | CLCF1 | 0.791 |
| | DUSP6 | -0.722 |
| | S1PR1 | -2.048 |
| | PTGS2 | -1.380 |

**Supplementary Table 2** Number of genes significantly changed in each treatment when compared to either the untreated sample or the time-matched, media control.

| Treatment | Compared to: | Number of Genes Changed |
|---|---|---|
| IMDM | untreated | 938 |
| TGF-β1 | untreated | 1085 |
| | IMDM | 117 |
| DMEM/F12 | untreated | 2879 |
| M4 TCM | untreated | 2740 |
| | DMEM/F12 | 27 |

### Figure S1

dHL-60 cells

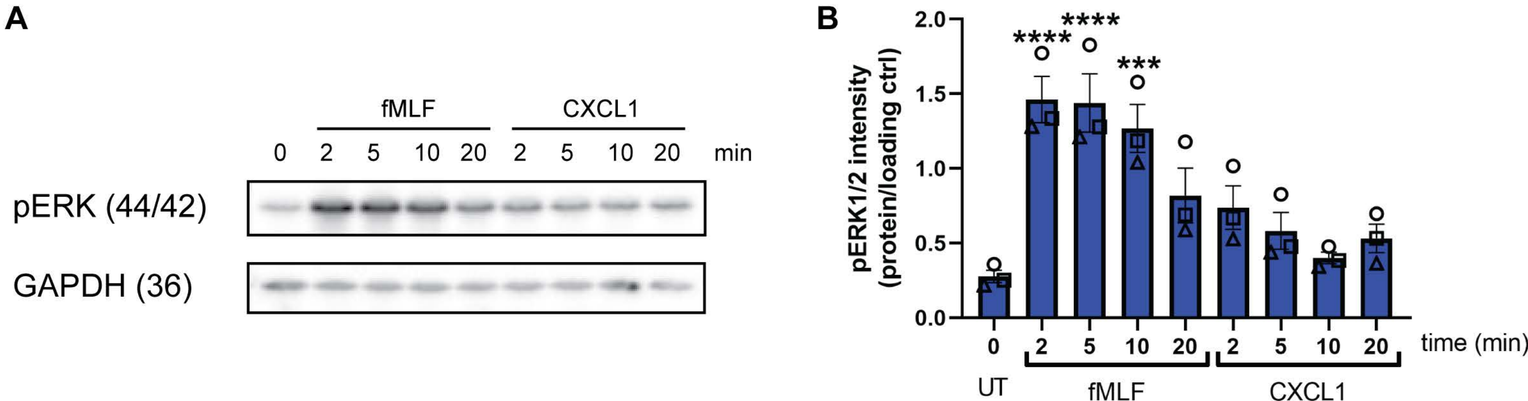

### Figure S2

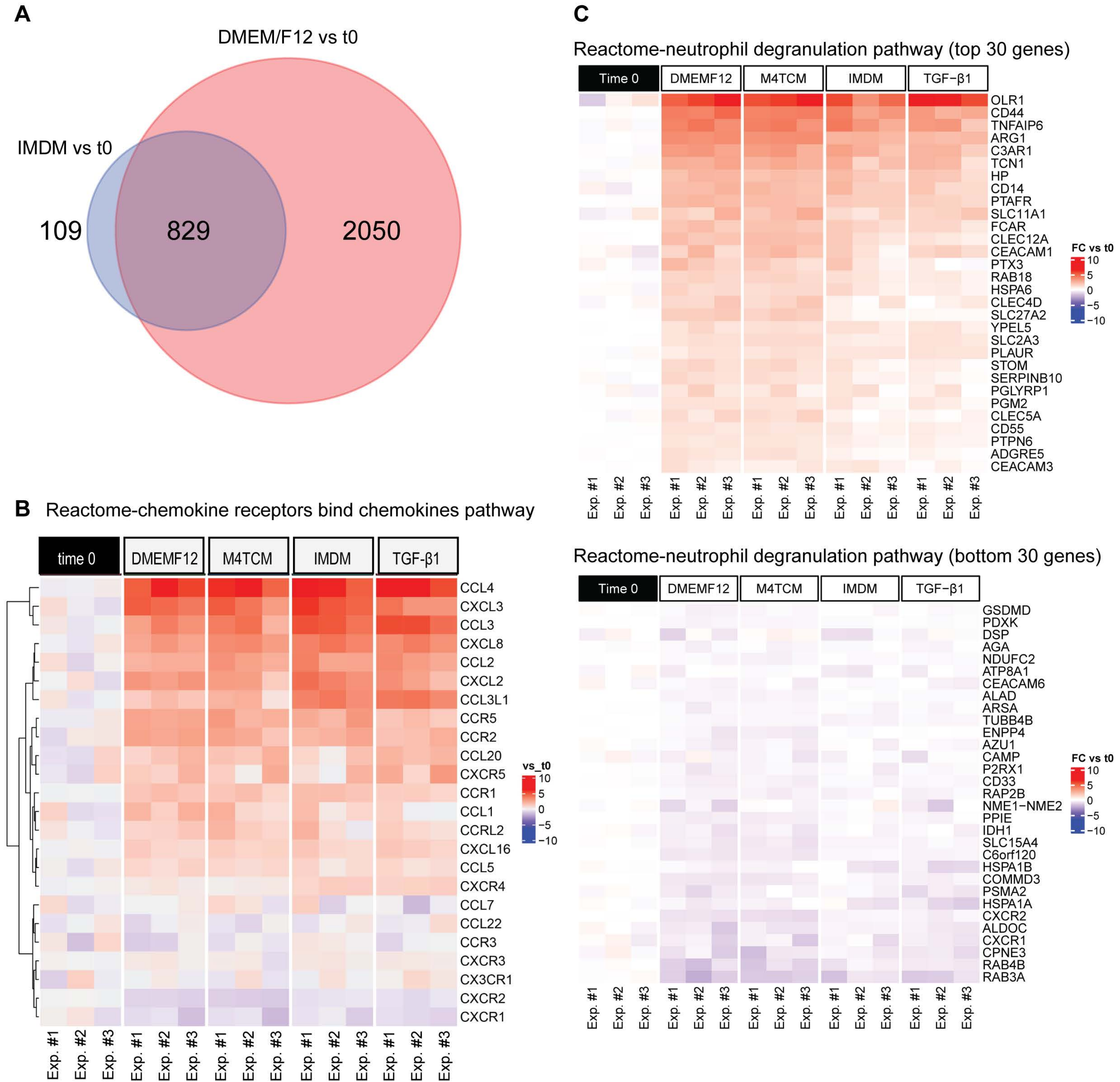
A
DMEM/F12 vs t0
IMDM vs t0
109
829
2050
B
Reactome-chemokine receptors bind chemokines pathway
time 0
DMEMF12
M4TCM
IMDM
TGF-β1
CCL4
CXCL3
CCL3
CXCL8
CCL2
CXCL2
CCL3L1
CCR5
CCR2
CCL20
CXCR5
CCR1
CCL1
CCRL2
CXCL16
CCL5
CXCR4
CCL7
CCL22
CCR3
CXCR3
CX3CR1
CXCR2
CXCR1
vs_t0
10
5
0
−5
−10
Exp. #1
Exp. #2
Exp. #3
C
Reactome-neutrophil degranulation pathway (top 30 genes)
Time 0
DMEMF12
M4TCM
IMDM
TGF-β1
OLR1
CD44
TNFAIP6
ARG1
C3AR1
TCN1
HP
CD14
PTAFR
SLC11A1
FCAR
CLEC12A
CEACAM1
PTX3
RAB18
HSPA6
CLEC4D
SLC27A2
YPEL5
SLC2A3
PLAUR
STOM
SERPINB10
PGLYRP1
PGM2
CLEC5A
CD55
PTPN6
ADGRE5
CEACAM3
FC vs t0
10
5
0
−5
−10
Exp. #1
Exp. #2
Exp. #3
Reactome-neutrophil degranulation pathway (bottom 30 genes)
Time 0
DMEMF12
M4TCM
IMDM
TGF-β1
GSDMD
PDXK
DSP
AGA
NDUFC2
ATP8A1
CEACAM6
ALAD
ARSA
TUBB4B
ENPP4
AZU1
CAMP
P2RX1
CD33
RAP2B
NME1−NME2
PPIE
IDH1
SLC15A4
C6orf120
HSPA1B
COMMD3
PSMA2
HSPA1A
CXCR2
ALDOC
CXCR1
CPNE3
RAB4B
RAB3A
FC vs t0
10
5
0
−5
−10
Exp. #1
Exp. #2
Exp. #3
